# The cannabinoid agonist CB-13 produces peripherally mediated analgesia in mice but elicits tolerance and signs of CNS activity with repeated dosing

**DOI:** 10.1101/2021.04.23.441212

**Authors:** Richard A. Slivicki, Jiwon Yi, Victoria E. Brings, Phuong Nhu Huynh, Robert W. Gereau

## Abstract

Activation of cannabinoid receptor type 1 (CB_1_) produces analgesia in a variety of preclinical models of pain; however, engagement of central CB_1_ receptors is accompanied by unwanted side effects, such as tolerance and dependence. Efforts to develop novel analgesics have focused on targeting peripheral CB_1_ receptors to circumvent central CB_1_-related side effects. In the present study, we evaluated the effects of acute and repeated dosing with the peripherally selective CB_1_-preferring agonist CB-13 on nociception and central CB_1_-related phenotypes in an inflammatory model of pain in mice. We also evaluated cellular mechanisms underlying CB-13-induced antinociception *in vitro* using cultured mouse dorsal root ganglion (DRG) neurons. CB-13 reduced inflammation-induced mechanical allodynia in a peripheral CB_1_ receptor-dependent manner and relieved inflammatory thermal hyperalgesia. In cultured mouse DRG neurons, CB-13 reduced TRPV1 sensitization and neuronal hyperexcitability induced by the inflammatory mediator prostaglandin E2, providing potential mechanistic explanations for the analgesic actions of peripheral CB_1_ receptor activation. With acute dosing, phenotypes associated with central CB_1_ receptor activation occurred only at a dose of CB-13 approximately 10-fold the ED_50_ for reducing allodynia. Strikingly, repeated dosing resulted in both analgesic tolerance and CB_1_ receptor dependence, even at a dose that did not produce central CB_1_ receptor-mediated phenotypes on acute dosing. This suggests repeated CB-13 dosing leads to increased CNS exposure and unwanted engagement of central CB_1_ receptors. Thus, caution is warranted regarding therapeutic use of CB-13 with the goal of avoiding CNS side effects. Nonetheless, the clear analgesic effect of acute peripheral CB_1_ receptor activation suggests that peripherally restricted cannabinoids are a viable target for novel analgesic development.

## Introduction

Chronic pain impacts 11.2% of individuals nationwide[53] and current pharmacotherapies for the treatment of chronic pain produce significant unfavorable effects including tolerance and dependence. The U.S. is currently in the midst of an opioid epidemic that is driven in part by opioids prescribed for pain relief [37]. Thus, there is an urgent need to develop safer, more effective therapies for the treatment of chronic pain. A large body of preclinical evidence supports the use of cannabinoids as pain-relieving pharmacotherapies in rodent models of neuropathic, inflammatory and visceral pain [2,77]. This mirrors some clinical reports showing vaporized or smoked cannabis is effective at treating different types of chronic pain [31]. However, a recent IASP task force [1] concluded that the current evidence base is insufficient to endorse the use of cannabis for the clinical management of pain, and called for “more rigorous and robust research to better understand any potential benefits and harms.” Cannabis-based therapies are not always efficacious and can possess unwanted side effects, including tolerance and psychoactivity [55]. Such centrally mediated side effects may limit therapeutic use by prevent adequate dosing to produce a peripherally mediated analgesic effect. Furthermore, it is important to note that Δ^9^-THC, the primary active ingredient in the cannabis-based therapies tested in clinical trials, is a partial agonist at CB_1_ receptors [59,67]. Thus, it possible that full activation of peripheral CB_1_ receptors could produce more robust analgesia in clinical trials than cannabis-based treatments have shown to date while also avoiding centrally mediated side effects. Therefore, considerable interest remains in developing peripherally restricted cannabinoid receptor agonists for pain and other indications, building on prior evidence supporting the use of cannabis-based medicines for the treatment of pain.

Cannabinoids exert their effects through engagement of cannabinoid receptors, which include cannabinoid type-1 (CB_1_) and type-2 (CB_2_) receptors. In particular, CB_1_ receptor engagement has been demonstrated to reduce pain-related behaviors in a number of animal models [77]. Activation of CB_1_ receptors in the central nervous system also results in psychoactivity in humans and typical “tetrad” behaviors in rodents (catalepsy, hypothermia, motor ataxia and antinociception[44]). Thus, utilizing CB_1_ receptor signaling in a manner that provides analgesia while avoiding such side effects would be ideal for therapeutic use.

There are many potential sites of analgesic action for CB_1_ receptor activation, as these receptors are expressed abundantly throughout the central and peripheral nervous systems[30,32,33,77]. Many studies show analgesic effects of peripheral CB_1_ activation. Peripheral administration of cannabinoid agonists leads to reductions of neuropathic allodynia[22,52], and inhibition of the endocannabinoid degradative enzyme fatty-acid amide hydrolase in the periphery results in peripheral CB_1_ receptor-mediated reductions in nociceptive-like behaviors in inflammatory and neuropathic pain models [7,68]. Deletion of CB_1_ receptors from primary sensory neurons leads to a loss of antinociceptive efficacy of WIN55,212-2 (a centrally penetrant pan-cannabinoid agonist [11]), while typical behaviors reflective of central CB_1_ engagement (e.g. catalepsy) are conserved [3]. Thus, targeting peripheral CB_1_ receptors may conserve analgesic efficacy while circumventing unwanted side effects associated with engagement of central CB_1_ receptors. Drug discovery efforts have therefore focused on designing peripherally restricted CB_1_ receptor-preferring ligands. Indeed, peripherally restricted cannabinoid agonists reduce nociceptive-like behaviors in models of neuropathic [22,81,83], inflammatory [21,81] and headache [78] pain when administered acutely. Many of the aforementioned studies, however, do not include pharmacological or genetic control conditions to ensure the analgesic effects of such compounds are mediated by peripheral CB_1_ receptors. Further, consequences of long-term, repeated administration of such compounds on both centrally mediated behaviors and antinociception remain poorly understood.

The present study sought to evaluate the peripherally selective CB_1_ receptor-preferring agonist CB-13[22] in both male and female mice using a model of inflammatory pain induced by Freund’s complete adjuvant (CFA), as this model has previously been shown to be unresponsive to CB_2_ agonists in mice [41]. We also assessed the impact of repeated administration of CB-13 on therapeutic tolerance and the cannabinoid ‘triad’ (catalepsy, antinociception, hypothermia) in CFA-treated and naïve animals, respectively. Finally, we evaluated the ability of CB-13 to reduce prostaglandin E2 (PGE2)-induced TRPV1 sensitization and alterations in neuronal excitability as potential mechanistic links to CB-13’s analgesic efficacy.

## Methods

### Subjects

All experiments used C57BL/6J mice purchased from Jackson Laboratory (Bar Harbor, ME) or bred in-house. Mice were 8-9 weeks of age at the start of behavioral experiments and 6-8 weeks of age for calcium imaging and electrophysiology experiments [66]. Animals were group housed with 3-5 animals per cage and were maintained in a temperature-controlled facility with *ad libitum* food and water and maintained on a 12-hour light-dark cycle (lights on at 07:00 hr – 19:00 hr). All experimental procedures were approved by the Washington University Animal Care and Use Committee and followed the guidelines of the Internal Association for the Study of Pain. Mice were randomly assigned to experimental conditions.

### Drugs and chemicals

Freund’s complete adjuvant (CFA) (Thermo Fisher, St. Louis, MO) was dissolved in a 1:1 ratio of saline:CFA prior to intraplantar (i.pl) injection. CB-13, AM6545 and rimonabant (all from Cayman Chemical Company, Ann Arbor, MI) were dissolved in vehicle consisting of 20% DMSO (Sigma Aldrich, St. Louis, MO), 8% ethanol, 8% 1Tween 80 (Thermo Fisher, St. Louis, MO) and 64% saline and administered via intraperitoneal (i.p.) injection in a volume of 5 mL/kg for behavioral studies. CB-13 and rimonabant were dissolved in DMSO (25 mg/mL) and frozen until use. On the day of the experiment, stocks were diluted with a volume of DMSO to achieve a concentration of 20% DMSO in the final solution. 95% ethanol was then added, the solution vortexed again, followed by Tween 80 and finally saline. For AM6545, the steps were the same as the other compounds used, but the solution was sonicated for approximately 45 minutes to get the compound to fully dissolve (when the solution was clear). For calcium imaging and electrophysiology studies, prostaglandin E2 (PGE2) (Thermo Fisher, St. Louis, MO) and CB-13 were dissolved in DMSO and diluted in external recording solution.

### Allodynia measurements

Mechanical allodynia was evaluated using an electronic von Frey anesthesiometer (IITC Life Science, model Alemo 2390–5, Woodland Hills, CA) as described previously [19,28,68,69]. In brief, animals were habituated to 10 cm x 10 cm acrylic holding containers placed on an elevated mesh platform 1 h prior to testing. Black dividers prevented mice from seeing other mice being tested simultaneously. Pressure was applied to the plantar surface of the hindpaw using the anesthesiometer, and the maximum force applied before the animal withdrew its paw was recorded. Each paw was stimulated twice with at least 7 min between stimulations. The two withdrawal thresholds were averaged for each data point.

Responsivity to heat stimulation was measured using a thermal plantar test apparatus (IITC Life Science, model 390) as described previously[5]. In brief, animals were habituated on a glass surface heated to 30°C in the same acrylic containers with black dividers as used for mechanical allodynia. A focused beam of light was applied to the hindpaw, and the latency for the animal to withdraw its paw was recorded. Each hindpaw of the animal was tested 2-3 times with at least 5 min between each measurement. A cutoff latency of 20 s per stimulation was applied to avoid tissue damage. The withdrawal latencies for all stimulations on a single paw were averaged for each data point.

### CFA-induced allodynia

CFA has been widely used to model allodynic responses following an inflammatory insult [48]. To induce CFA-induced injury, on the same day as baseline values were recorded, a 20 μL injection of 1:1 CFA:saline was administered via unilateral i.pl. injection in the hindpaw with a 28.5-gauge needle[69]. Approximately 18 h following CFA injection, post-CFA values for responsiveness to mechanical and heat stimulation were assessed.

### Dose-response studies

Within-subjects dose-response curves were generated to evaluate the ability of CB-13 to reduce CFA-induced mechanical allodynia as described previously[68,69]. CB-13 (0.3, 1, 3, 10 mg/kg i.p.) or vehicle was administered in spate cohorts of male and female mice starting approximately 20 h after CFA injection allowing at least 24 h between escalating doses. Mechanical thresholds were evaluated prior to CB-13 or vehicle injection and 30 min following injection of the assigned drug condition.

### Timecourse and chronic dosing studies

In hourly timecourse experiments, animals were evaluated for mechanical or heat hypersensitivity (using separate cohorts for each modality) at baseline and 18 h following CFA injection. Immediately after post-CFA thresholds were measured, animals were injected with either vehicle or CB-13 (1, 3, 10 mg/kg i.p.) and then evaluated again for responsivity to mechanical (0, 0.5, 1, 1.5, 6, 7.5, and 24 h) or heat (0, 0.5, 1, 1.5, 2, 5, 7.5, and 24 h) stimulation. Doses were chosen based on dose-response response studies and previous reports. A low but effective dose (1 mg/kg i.p.) [8], a maximally effective but peripherally restricted dose (3 mg/kg i.p.) [60,61] and a high but likely CNS-penetrant dose (10 mg/kg i.p.) [60] were incorporated to better understand the interplay between peripheral and central activation of CB_1_ receptors in relation to anti-allodynic effects and tolerance development. For chronic dosing studies, the same animals tested in the mechanical hypersensitivity hourly timecourse were administered the assigned treatment once daily at ∼10:00 h. Animals were evaluated for mechanical thresholds prior to and 30 min post injection on days 1, 3 and 7. Chronic dosing was not evaluated for heat hypersensitivity as our studies indicated that thresholds returned to baseline values ∼26 h after CFA injection (Figure 4E).

### Pharmacological specificity

In CFA-injected animals, the peripherally restricted CB_1_ receptor antagonist AM6545 (10 mg/kg i.p.)[9] was administered 30 min prior to administration of CB-13 (3 mg/kg i.p.), or 60 min prior to testing. Animals were tested at 30 min following CB-13 injection, as this is the timepoint at which the reversal of allodynia was deemed to be maximal based on the timecourse studies (Figure 1E).

**Figure 1:**
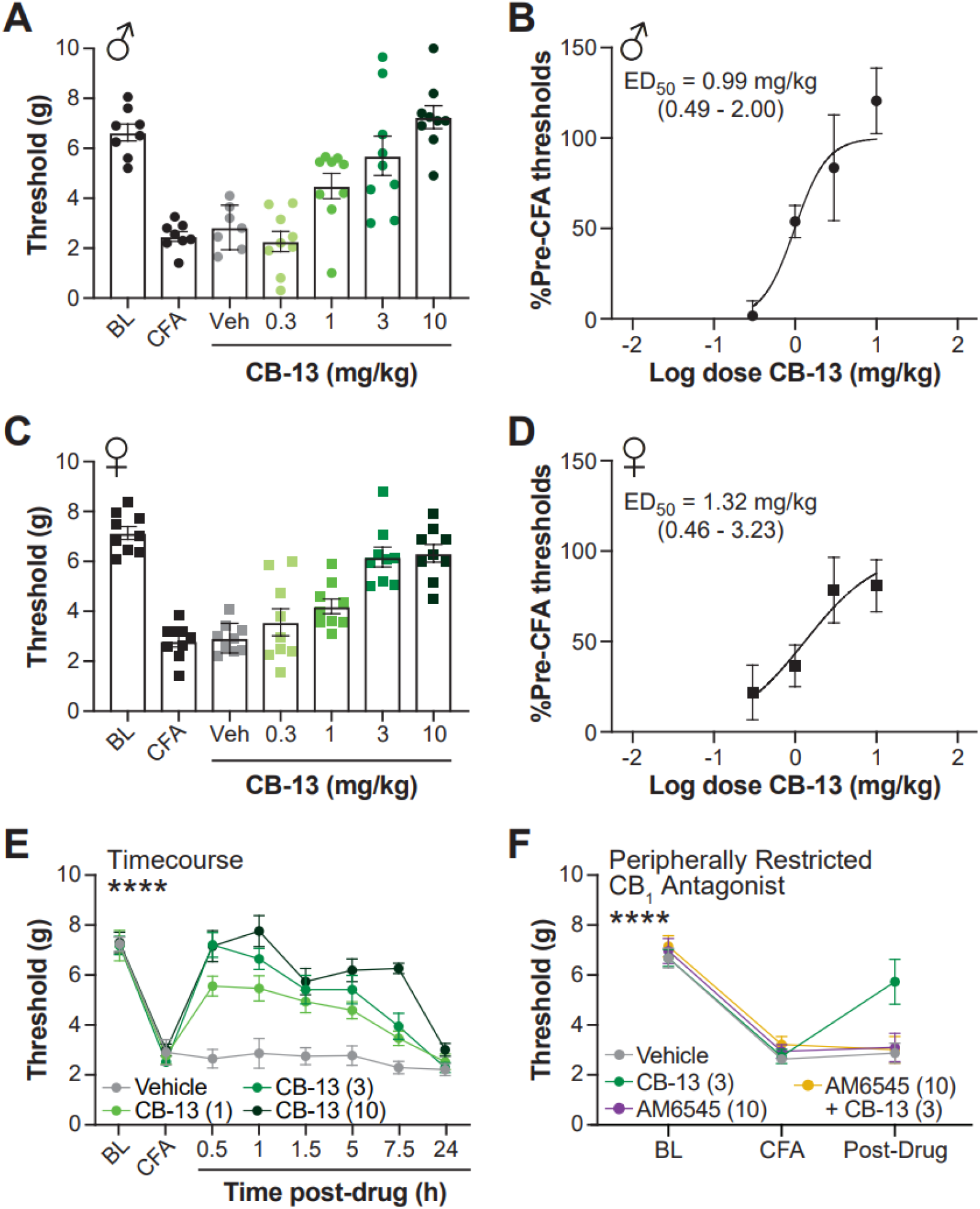
Administration of CB-13 produces a long-lasting dose-dependent reduction in CFA-induced mechanical allodynia that is dependent on CB_1_ receptor activation in the periphery. CB-13 (0.3, 1, 3, 10 mg/kg i.p.) reduced CFA-induced mechanical allodynia in a dose-dependent manner that was equipotent in both males (A,B) and females (C,D). CB-13’s anti-allodynic effects are long-lasting (E) (BL = baseline, CFA = 24hr post-CFA, but pretreatment with vehicle or CB-13). Pre-treatment with the peripherally restricted CB_1_ receptor antagonist AM6545 (10 mg/kg i.p.) prevents the analgesic effect of CB-13 (3 mg/kg i.p.), but has no effect on its own (F). ED_50_ values (in mg/kg) displayed with 95% confidence intervals on graphs in (B) and (D). **** P < 0.0001 2×2 ANOVA interaction effect all timepoints post-CFA. (E). **** P < 0.0001 One-way ANOVA post-drug timepoint (F). Post-tests and other statistics are reported in Table 1. Data are expressed as mean ± SEM. N = 7-9 per group.

### Cannabinoid triad

Centrally penetrant CB_1_ receptor agonists induce a classical “tetrad” of behavioral effects in mice which include catalepsy, locomotor ataxia, antinociception and hypothermia[44]. We examined, in order, catalepsy, tail-flick antinociception and hypothermia at several different timepoints. Animals were dosed once daily with either vehicle or CB-13 (3 or 10 mg/kg i.p.). Triad measures were taken at several times (just prior to dosing and 0.5, 1, 1.5, 3, 6, 7.5 and 24 h post-drug) on days 1, 3 and 7 to mimic chronic dosing studies in the CFA condition.

### Evaluation of CB_1_ receptor-mediated dependence

CB_1_ receptor dependence is commonly evaluated via administration of the CB_1_ antagonist rimonabant to precipitate somatic withdrawal in animals chronically treated with a CB_1_ agonist[42,43]. In chronic dosing studies, mice were dosed daily with CB-13 or vehicle for 9 days, and on the 9^th^ day rimonabant (10 mg/kg i.p.) was administered 1 h following a final injection of CB-13 or vehicle, and animals were video recorded for 30 min. The number of rimonabant-evoked scratching behaviors and paw tremors were manually evaluated using Behavioral Observation Research Interactive Software (BORIS)[24].

### Mouse DRG cultures

For each tissue preparation, mice were euthanized via induction of isoflurane briefly followed by decapitation. The spinal column was removed and bisected; lumbar DRG were removed and pooled, and subjected to enzymatic incubations as described [16,45,66]. DRG tissue was incubated in papain (45 U in HBSS+H, Worthington) for 20 min at 37°C, rinsed, and incubated in collagenase (1.5 mg/mL in HBSS+H, Sigma-Aldrich) for 20 min. DRG were washed again with HBSS+H, then transferred to 1mL of DRG media which consisted of Neurobasal A medium (Invitrogen) supplemented with 100 U/mL penicillin/streptomycin (Corning), 2 mM GlutaMAX (Life Technologies), 2% B27 (Gibco), and 5% fetal bovine serum (Gibco). DRG were then manually triturated with fire-polished Pasteur pipettes (VWR) and passed through a 40-μm filter (VWR). DRG were then plated onto poly-d-lysine/collagen (Sigma-Aldrich)-coated 12-mm glass coverslips (Thermo Fisher Scientific) and stored at 37°C and 5% CO_2_ until testing.

### Calcium imaging

Calcium imaging studies were performed the day after plating. Cultured neurons were incubated with 3 μg/mL of the ratiometric calcium indicator Fura-2 AM (Life Technologies) in external recording solution for 30 min at 37°C and 5% CO_2_. To evaluate the impact of CB_1_ receptor activation on TRPV1 sensitization, we utilized a protocol that assesses the ability of inflammatory mediators to potentiate capsaicin-induced calcium influx in cultured DRG neurons, as demonstrated and described in previous publications [66]. Coverslips were then transferred to a recording chamber and continuously perfused at room temperature with external recording solution, containing the following (in mM): 130 NaCl, 5 KCl, 2 CaCl_2_, 1 MgCl_2_, 30 glucose, and 10 HEPES. Cells were visualized under an inverted microscope (Olympus Optical), and fluorescent images were acquired every 2 s using a Hamamatsu ORCA camera (Hamamatsu) and SimplePCI Software (HCImage, Hamamatsu). Fura-2 AM fluorescence intensity was recorded using alternating excitation wavelengths of 357 and 380 nm. The experimental timeline was as follows: 2 min baseline (external solution), 20s application of 200 nM of capsaicin, 3 min wash with external solution, 11 min of drug treatment (1 of 3 conditions, follows), a second pulse of 200 nM of capsaicin was applied for 20s followed by a 4 min wash (external solution), then a 10s pulse of KCl (50 mM) was applied to evaluate for cell viability. The three drug treatment conditions were 11 min of vehicle, 1 min of vehicle followed by 10 min of 1 µM PGE2, or 1 min of 1 µM CB-13 followed by 10 min of 1 µM PGE2 + 1 µM CB-13. Each coverslip was tested with one treatment condition, and 2-3 treatment conditions were evaluated for each mouse preparation. Cells that did not return to within 10% of their baseline values after the first capsaicin application, or did not respond to KCl application, were excluded from further analysis.

### Electrophysiology

Electrophysiology experiments were conducted 1-2 days following plating. Primary DRG neurons were transferred to a recording chamber and perfused with external recording solution containing the following (in mM): 145 NaCl, 3 KCl, 2 CaCl_2_, 1 MgCl_2_, 7 glucose, 10 HEPES. Whole-cell patch-clamp recordings were made using fire-polished borosilicate glass pipettes with 3-6 MΩ resistance. The pipettes were filled with KCl-based internal solution consisting of the following (in mM): 120 K-gluconate, 5 NaCl, 3 MgCl_2_, 0.1 CaCl_2_, 10 HEPES, 1.1 EGTA, 4 Na_2_ATP, 0.4 Na_2_GTP, 15 Na_2_Phosphocreatine; adjusted to pH 7.3 with KOH and HCl, and 290 mOsm with sucrose. For some of the recordings, the internal solution also contained 50 µM Alexa Fluor 568 (Invitrogen). All experiments were conducted at room temperature. Recordings were made using a MultiClamp 800B amplifier and a Digidata 1550B digitizer (Molecular Devices, CA). Only smaller neurons with a diameter <24 µm were studied (19.0 ± 2.3 µm, average ± SD). Series resistance was kept below 15 MΩ in all recordings. After a stable whole-cell configuration was achieved, membrane excitability was assessed in current-clamp mode. All recordings were done at the resting membrane potential of each cell (−53.6 ± 7.1 mV at baseline, average ± SD). For all experiments, membrane excitability was assessed before (“pre-treatment”) and after drug treatment (1 μM PGE2, 1 μM CB-13, 1 μM CB-13 + 1 μM PGE2, or 0.1% DMSO). In a subset of CB-13 experiments (n = 5 cells), cells were additionally treated with both the CB-13 + PGE2 condition following the CB-13 incubation. There was no significant difference in membrane excitability during CB-13 + PGE2 treatment between cells that were pre-treated with CB-13 and cells that were not.

Resting membrane potential was monitored during drug application and throughout the experiment; if any cell’s resting membrane potential depolarized >-40 mV, the experiment was discarded or excluded from analysis. Electrophysiology data were compiled using ClampFit (v.11.1, Molecular Devices). Input-output relationship was determined by counting the number of action potentials generated by 1 second current steps in 10 pA increments from 0 to 300 pA. To account for the heterogeneity in firing patterns between cells, the input-output curve was normalized to each cell’s current threshold at its pre-treatment baseline. The current threshold was established from the current step at which the first action potential was generated. Input resistance was determined as ΔV/ΔI using 10-50 pA hyperpolarizing current injections. First action potential elicited at rheobase was used to measure the action potential threshold, defined as the voltage at which the slope of the action potential (dv/dt) equals 10.

### Statistical analyses

The experimenter (RS) was blind to behavioral treatment condition and throughout analysis of calcium imaging data. All data were analyzed using Graphpad Prism 8.0 and Excel. Raw data for ED_50_ calculations were converted to % baseline responding (i.e. prior to CFA treatment) using the following formula: (Experimental Value – Post-CFA baseline)/(Pre-CFA baseline – post-CFA baseline). ED_50_ values were generated using nonlinear regression analysis in GraphPad 8.0. Behavioral data were analyzed via an ANOVA (two-way or one-way) followed by Tukey’s post-hoc. In the case of only two groups, Bonferroni’s post-hoc was used (Figure 4A). A one-way ANOVA followed by Tukey’s post-hoc was used to compare the response ratio for calcium imaging groups. For all electrophysiology datasets, normality of residuals was tested using the Kolmogorov-Smirnov test, and parametric and non-parametric tests were used accordingly. Changes in current threshold and input resistance post-drug across conditions were analyzed using one-way ANOVA with Tukey’s post-hoc. Change in voltage threshold post-drug across conditions was analyzed using Kruskal-Wallis test. Difference between pre- and post-drug current threshold and input resistance for each drug condition was evaluated using paired t-tests (parametric) and Wilcoxon matched-pairs signed rank test (non-parametric). Resting membrane potential across time was analyzed using one-way repeated measures ANOVA. For input-output data, mixed effects model with Tukey’s post-hoc test was used for analysis. Detailed statistics including main effects and post-hoc comparisons are reported in the tables.

## Results

### CB-13 produces long-lasting analgesia mediated by peripheral CB_1_ receptors in male and female mice

CFA induced mechanical allodynia in both male and female mice, and this allodynia was dose-dependently reduced by the peripherally restricted cannabinoid agonist CB-13, administered at approximately 20 h post-CFA injection. The ED_50_ for reduction of CFA-induced mechanical allodynia by CB-13 was 0.99 mg/kg (95% CI 0.49 -2.00 mg/kg) in male mice (Figure 1A,B) and 1.32 mg/kg (95% CI 0.46 -3.23 mg/kg) in female mice (Figure 1C,D) when measured 30 min post-drug injection. The ED_50_ values are comparable and the CIs overlap, suggesting efficacy of CB-13 did not differ between sexes. Male mice were used for all subsequent experiments. To determine the timecourse of CB-13 anti-allodynia, CB-13 (1, 3, or 10 mg/kg i.p.) or vehicle was administered after confirming CFA-induced hypersensitivity, as described above, and mechanical sensitivity was assessed for 24 h post-drug delivery. The reduction in CFA-induced mechanical allodynia by CB-13 was long-lasting (Fig. 1E; F_18, 150_ = 5.391, p<0.0001, two-way ANOVA interaction effect). This effect appeared to be maximal at 30 min post-injection and lasted for at least 6 h for each dose (Figure 1E). The anti-allodynic effect of CB-13 (3 mg/kg i.p.) was completely abolished by the peripherally restricted CB_1_ antagonist AM6545 (10 mg/kg i.p., administered 30 min prior to CB-13). Administration of AM6545 did not alter mechanical thresholds on its own (Table 1; Figure 1F). These results demonstrate that the analgesic effects of CB-13 are mediated exclusively by peripheral CB_1_ receptors.

### Repeated administration with CB-13 results in antinociceptive tolerance and CB_1_ receptor-mediated dependence

We next evaluated if CB-13 maintained efficacy over a repeated dosing schedule, or if tolerance would develop. CB-13 (1, 3, or 10 mg/kg i.p.) reduced CFA-induced mechanical allodynia (F_9, 72_= 7.792, p<0.0001, two-way ANOVA interaction effect) on days 1 and 3 of chronic daily dosing (Figure 2A; Table 2). By day 7 of repeated dosing, analgesic tolerance appeared to develop to all doses of CB-13.

**Figure 2.**
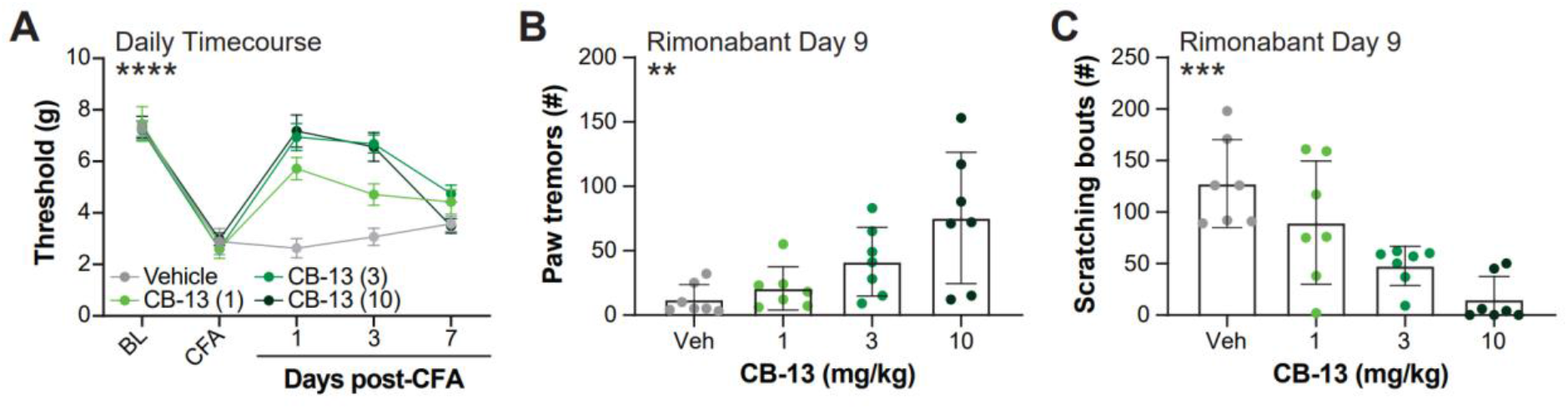
Sustained administration of CB-13 produces analgesic tolerance and CB_1_ receptor dependence in CFA-treated mice. Following baseline (BL) assessment and the induction of CFA-induced inflammation, animals were randomized to receive 1, 3 or 10 mg/kg i.p. of CB-13 or vehicle and evaluated for mechanical responsivity 30 min following treatment administration on days 1, 3 and 7. All doses were effective on days 1 and 3, but by day 7 tolerance developed to the anti-allodynic effect of CB-13 (A). On day 9 the CB_1_ antagonist rimonabant (10 mg/kg i.p.) was administered 1 h following the final injection of CB-13 or vehicle. Chronic treatment with 3 or 10 mg/kg increased rimonabant-elicited paw tremors (B) and decreased rimonabant-elicited scratching behaviors (C), consistent with the development of CB_1_ receptor dependence. ****P<0.0001 2×2 ANOVA interaction effect in (A). ** P < 0.01, *** P < 0.001 One-way ANOVA effect of treatment in (B) and (C). Post-tests and other statistics are reported in Table S2. Data are expressed as mean ± SEM. N = 7-9 per group.

The development of analgesic tolerance to CB_1_ receptor agonists is typically concurrent with CB_1_-mediated dependence [18,39]; therefore, we sought to evaluate if the animals receiving repeated doses of CB-13 would also demonstrate CB_1_-mediated dependence. On day 9 of daily dosing, animals received a final injection of the assigned treatment condition, followed by administration of the CB_1_ antagonist rimonabant (10 mg/kg i.p.) 1 h later. In animals treated with CB-13, rimonabant administration elicited increased paw tremors (F_3, 24_ = 5.971, p=0.0034, one-way ANOVA; Figure 2B). Treatment with CB-13 also reduced rimonabant-evoked scratching behaviors (F_3,24_ = 10.78, p=0.0001, one-way ANOVA; Table 2; Figure 2C), consistent with the development of CB_1_ receptor dependence following repeated dosing with CB-13 even at doses that were previously reported[8] to be peripherally restricted and devoid of CNS side effects.

### CB-13 induces cardinal signs of central CB_1_ receptor activation independent of inflammatory status

The development of anti-allodynic tolerance and rimonabant-evoked dependence phenotypes suggests potential engagement of central CB_1_ receptors over repeated dosing. To determine the timepoints in the repeated dosing regimen at which central CB_1_-mediated phenotypes emerge, we evaluated CB-13’s effects on catalepsy, tail-flick antinociception and body temperature, which are phenotypes affected by central CB_1_ receptor activation [20,26,47], on days 1, 3 and 7 of daily CB-13 treatment in naïve (non-CFA-treated) mice. CB-13 increased the time spent immobile in the bar test of catalepsy behavior at the highest dose of 10 mg/kg i.p. on the first day of testing (F_14, 112_ = 8.983, p<0.0001, two-way ANOVA interaction effect), with the peak cataleptic effect occurring at 7.5 h post-injection (Figure 3A; Table 3). CB-13 also produced catalepsy on day 3 (F_14, 112_ = 3.347, p=0.0002, two-way ANOVA interaction effect; Figure 3B) but did not induce catalepsy by day 7 of repeated dosing (Figure 3C). Importantly, catalepsy did not occur until 6 h post-injection of 10 mg/kg CB-13 (Figure 3A); thus, because the effects of CB-13 on CFA-induced allodynia were measured at 0.5 h post-drug administration (Figure 1; Figure 2A) and were noted to be maximal at that timepoint (Figure 1E), catalepsy was not a confounding variable in the anti-allodynia observed. CB-13 induced tail-flick antinociception (F_14, 112_ = 5.921, p<0.0001, two-way ANOVA interaction effect) at the highest dose tested (10 mg/kg i.p.) at 3 h post-injection (Figure 3D; Table 3). This effect was no longer evident on day 3 or 7 of sustained dosing (Figure 3E-F), consistent with the development of tolerance. CB-13 did not induce significant alterations in body temperature at any timepoint as measured by post-hoc analysis, although a trend to hypothermia is observed on day 1 at the highest dose tested (F_14,112_ =2.255, p=0.0096, two-way ANOVA interaction effect; Figure 3G-I). Collectively, these data are consistent with the engagement of central CB_1_ receptors at a CB-13 dose of 10 mg/kg, even with acute dosing.

**Figure 3.**
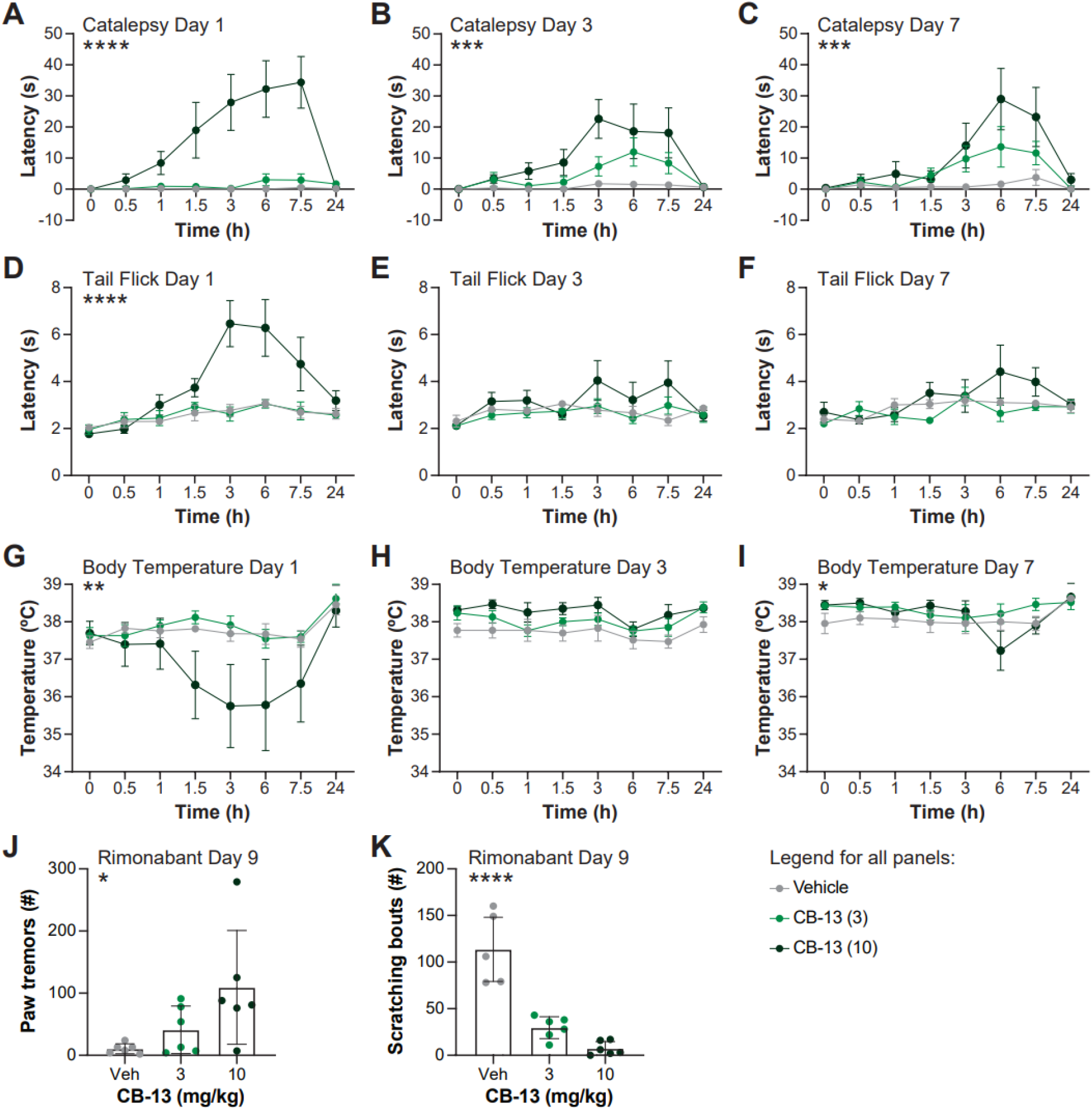
CB-13 produces cardinal signs of CNS CB_1_ receptor activation and CB_1_ receptor dependence in CFA-naïve mice. Mimicking the sustained treatment of Figure 2, we evaluated CB-13 for its potential to produce centrally mediated effects of CB_1_ receptor activation in non-CFA-treated mice. Otherwise naïve animals were administered CB-13 (3 and 10 mg/kg i.p.) or its vehicle and evaluated (in order) for catalepsy, tail-flick antinociception and hypothermia. The highest dose of CB-13 produced catalepsy on day 1 (A); however, by day 3 (B) and day 7 (C) this effect was no longer significant. The highest dose of CB-13 produced tail-flick antinociception on day 1 (D) but failed to do so on days 3 (E) and 7 (F), suggesting tolerance developed to this effect. Hypothermia was not induced on any day at any dose (G-I). Following a final injection of CB-13 on day 9, rimonabant (10 mg/kg i.p.) was administered to evaluate for the occurrence of CB_1_ receptor dependence. Similar to Figure 2, there was an increase in paw-tremor (J) and decrease in rimonabant-evoked scratching (K) behaviors. *P<0.05, **P < 0.01, *** P < 0.001, **** P < 0.0001 2×2 ANOVA interaction effect. *P<0.05, ****P < 0.0001 One-way ANOVA effect of treatment. Post-tests and other statistics are reported in Table S3. Data are expressed as mean ± SEM. N = 6-7 per group.

To evaluate if CB-13-induced CB_1_ dependence was impacted by an inflammatory state, we tested whether chronic administration of CB-13 in naïve animals not administered CFA would also result in CB_1_ receptor dependence. As with the CFA-treated mice (Figure 2B-C), rimonabant was administered on day 9 after a final dose of CB-13 (3 or 10 mg/kg i.p.) or vehicle in naïve mice. Chronic dosing with CB-13 again dose-dependently increased rimonabant-induced paw-tremors (F_2, 15_ = 4.637, p= 0.0271, one-way ANOVA; Figure 3J) and reduced rimonabant-elicited scratching behavior (F_2, 15_ = 40.94, p<0.0001, one-way ANOVA; Figure 3K), indicative of CB_1_ receptor-dependence (Table 3). This result suggests that CB_1_ receptor-dependence induced by CB-13 is not reliant on the presence of an inflammatory state.

### CB-13 reduces CFA-induced thermal allodynia

We next evaluated whether CB-13 can reduce thermal hyperalgesia following CFA administration, to test whether the analgesic effects of CB-13 would extend to other sensory modalities. CFA administration resulted in hypersensitivity to thermal stimulation (F_1,28_ = 109.1, p<0.0001, two-way ANOVA effect of time; Figure 4A). Using a dose that was maximally effective at reducing mechanical allodynia, CB-13 (3 mg/kg i.p.) reduced CFA-induced heat hypersensitivity (F_7, 196_ = 2.563, p=0.0151, two-way ANOVA interaction effect; Table 4).

**Figure 4.**
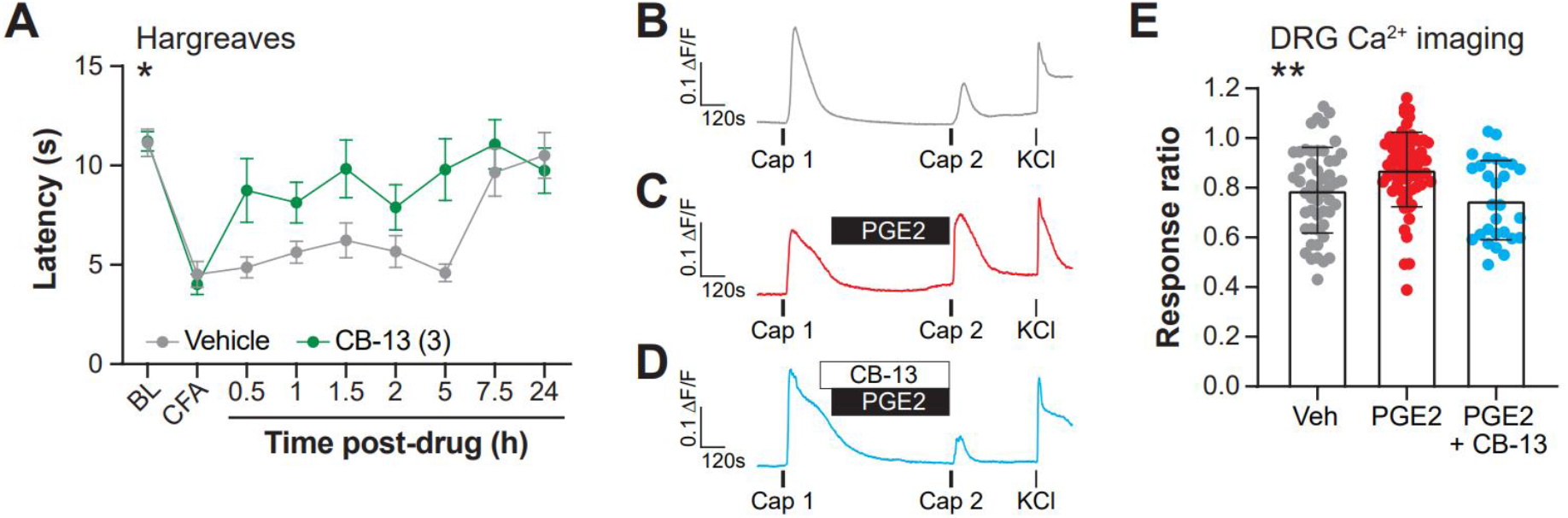
CB-13 reduces CFA-induced thermal allodynia and prevents PGE2-induced TRPV1 sensitization in mouse dorsal root ganglia neurons. CFA induces thermal allodynia (A, BL = baseline, CFA = 25 hr following CFA administration). CFA-induced thermal allodynia is reversed by administration of CB-13 (3 mg/kg, A). The dose used here was maximally effective at reducing CFA-induced mechanical allodynia and was previously reported to be peripherally restricted[60]. To evaluate a potential mechanistic link to this finding, we evaluated the effect of CB-13 on TRPV1 sensitization induced by PGE2 in mouse dorsal root ganglia neurons. Administration of capsaicin (200 nM) results in calcium influx in isolated mouse DRG neurons. A second identical application of capsaicin leads to a reduced, or desensitized response (B,E). This 2^nd^ capsaicin response is potentiated by application of the inflammatory mediator, PGE2 (1 μM, B,E). This potentiation of the 2^nd^ capsaicin response by PGE2 is prevented by co-application of CB-13 (1 μM; D,E). *P < 0.05 CB-13 vs. vehicle, 2×2 ANOVA interaction effect (A). **P < 0.01 One-way ANOVA. Post-tests and other statistics are reported in Table S4. Data are expressed as mean ± SEM for (A) and mean ± SD for (E). N = 15 mice per group (A), N = 28-63 cells from 4-8 mice per condition (E).

### CB-13 reduces PGE2-induced sensitization in cultured DRG neurons

We next sought to evaluate if CB-13 can reverse signs of peripheral sensitization induced by inflammatory mediators in isolated DRG neurons. We and others have shown that the inflammatory mediator PGE2 produces sensitization of the noxious heat transduction channel, TRPV1, and increases excitability in mouse DRG neurons [17,35,66]. We first assessed the ability of CB-13 to reduce PGE2-induced sensitization of TRPV1, as a plausible mechanism for reversal of inflammation-induced thermal hypersensitivity by CB-13 [34]. To this end, we employed calcium imaging of cultured mouse DRG neurons and confirmed the ability of PGE2 to sensitize responses to capsaicin application. Repeated application of capsaicin induced TRPV1 desensitization, as measured by a decrease in response to a second application of capsaicin (Figure 4B,E); when PGE2 was administered between capsaicin pulses, a sensitization effect was observed (Figure 4C,E; Table S4). When CB-13 (1 μM) was applied 1 min prior to PGE2 and co-applied during the duration of PGE2 administration, it prevented the PGE2-induced increase in the capsaicin response ratio (F_(2, 139)_ = 7.074, P=0.0012; Figure 4D-E; Table 4).

In addition to TRPV1 sensitization, PGE2 induces hyperexcitability of DRG neurons, which is another key peripheral process that contributes to nociceptive sensitization in the context of inflammation [4,16,51,72,79,80]. We used patch clamp electrophysiology to examine whether CB-13 can reduce PGE2-induced changes in DRG excitability. Bath application of 1 μM PGE2 significantly increased action potential firing in response to depolarizing current injections (F _(1.000, 10.00)_ = 11.82, P=0.0063, mixed effects model, PGE2 effect; Figure 5A, Table 5); PGE2-treated cells had significantly more spike firing compared to vehicle-treated cells (F _(3, 40)_ = 4.535, P=0.0079, mixed effects model, treatment effect; Figure 5E, Table 5). PGE2 additionally led to significant reduction in current threshold for action potential firing (F _(3, 40)_ = 7.534, P=0.0004, one-way ANOVA; Figure 5F; Table 5). The effect of PGE2 on spike firing was blocked by co-application of 1 μM CB-13 with PGE2 (Figure 5B,E; Table 5) but had no effect on PGE2-induced current threshold reduction. Application of CB-13 with PGE2 did not change the resting membrane potential, input resistance or voltage threshold for action potential of the recorded cell (Figure 5G-I; Table 5). To account for the potential effects of CB-13 or vehicle alone on action potential firing, we compared membrane excitability before and after treatment with either 1 μM CB-13 alone or vehicle. Neither CB-13 nor vehicle alone had any effect on the input-output curve, current threshold, input resistance, resting membrane potential or voltage threshold (Figure 5C-I; Table 5). Treatment with PGE2, PGE2+CB-13, CB-13 alone or vehicle did not lead to any changes in action potential firing phenotypes in response to depolarizing current injections. Together, these results suggest that CB-13 can partially attenuate PGE2-induced hyperexcitability in DRG neurons without affecting baseline membrane properties.

**Figure 5.**
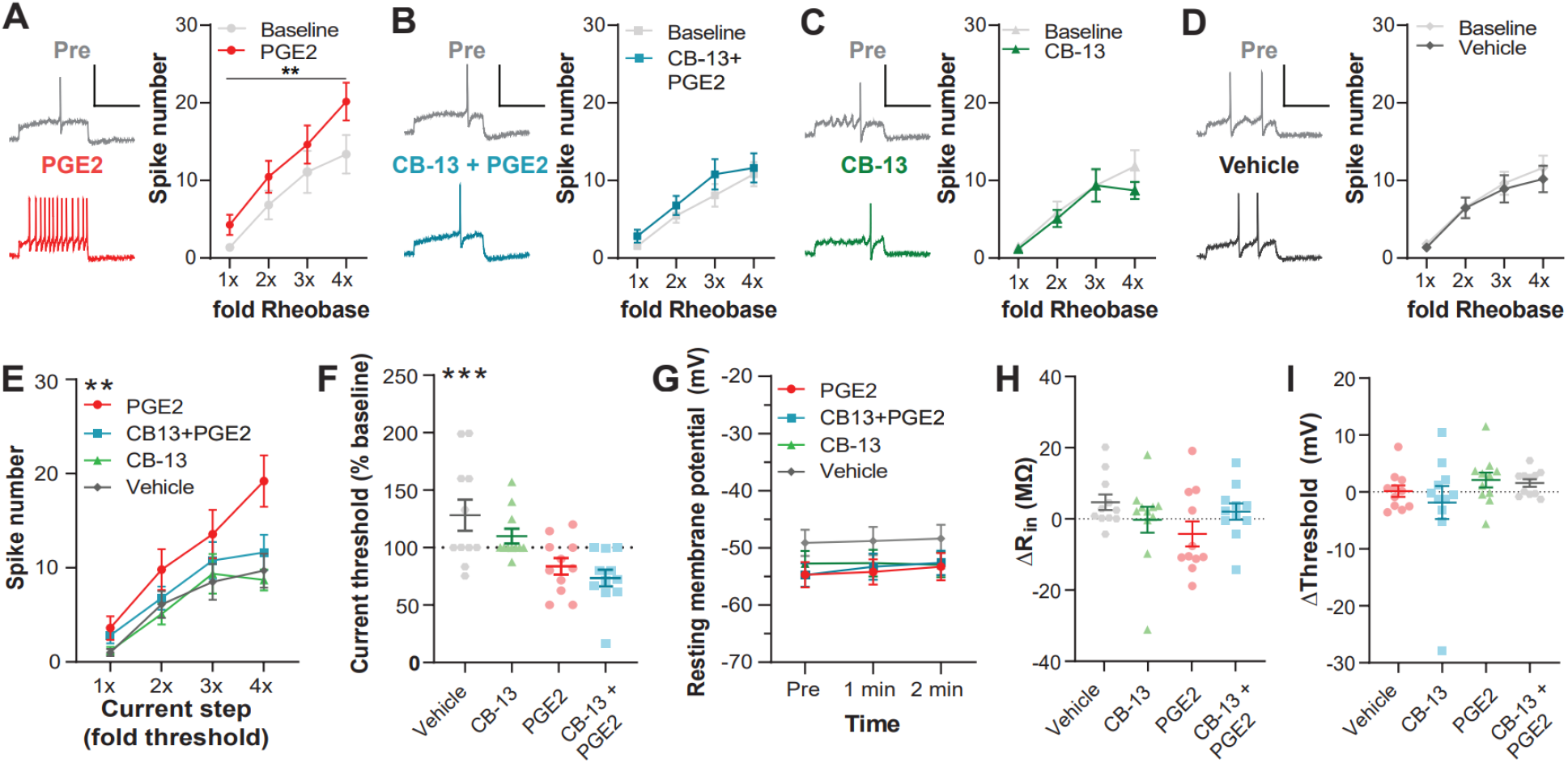
CB-13 attenuates PGE2-induced hyperexcitability of mouse DRG neurons *in vitro*. (A-D) Action potential firing before (grey) and after drug treatment: 1 µM PGE2 (A), 1 µM CB-13+PGE2 (B), CB-13 alone (C) or vehicle (0.1% DMSO; D). Right: Representative traces of action potentials at the minimum current threshold required to elicit an action potential (rheobase). Left: Input-output curve of number of action potentials evoked by depolarizing current injections, normalized to each cell’s current threshold. PGE2 treatment led to an upward shift in the input-output curve compared to baseline. This effect was not seen in cells treated with both CB-13 and PGE2. CB-13 or vehicle, when applied alone, did not induce any changes in spike firing. (E) Input-output curve post-treatment for all drug groups. There was a significant effect of PGE2 treatment on spike firing, which was blocked by CB-13 co-treatment. (F) CB-13 did not block PGE2-induced decrease in the minimum current required to elicit an action potential. CB-13 or PGE2 had no effect on resting membrane potential (G), input resistance (H) or voltage threshold for an action potential (I). ** P < 0.01; Mixed effects model, main effect of treatment (A, E). *** P < 0.001; One-way ANOVA followed by Tukey post-hoc (F). PGE2, N = 11 cells from 10 mice; CB-13+PGE2, N = 11 from 10 mice; CB-13, N = 11 from 9 mice; Vehicle, N = 11 from 8 mice. Post-tests and other statistics are reported in Table S5. Scale bars: 100 mV, 400 ms. Data are expressed as mean ± SEM.

## Discussion

In this study, we demonstrate that the peripherally selective CB_1_-preferring agonist CB-13 produces equipotent efficacy in reducing inflammation-induced allodynia in male and female mice. With acute administration, the analgesic effect of CB-13 is peripherally mediated and is achieved at doses that do not produce behaviors indicative of central CB_1_ receptor engagement. Surprisingly, however, behavioral indicators of central CB_1_ receptor activation are present after acute dosing at doses higher than required for anti-allodynia, or when dosing periods are extended over several days. Finally, CB-13 reduces measures of TRPV1 sensitization and cellular hyperexcitability induced by the inflammatory mediator PGE2 in cultured DRG neurons, supporting the notion that activating CB_1_ in DRG neurons can produce an analgesic effect, and providing a potential mechanistic basis for our behavioral findings on anti-allodynic effects of CB-13.

Centrally penetrant cannabinoids, such as Δ^9^-THC, exhibit differential effectiveness between sexes at reducing allodynia in models of inflammatory and neuropathic pain [10,12,13]. In contrast, the peripherally restricted cannabinoid agonist PrNMI showed similar degrees of efficacy between sexes in a rat model of chemotherapy-induced peripheral neuropathy [52]. Our study complements these results, as CB-13 produced dose-dependent reductions of CFA-induced mechanical allodynia with a similar degree of efficacy between sexes in mice. Importantly, acute anti-allodynic efficacy of CB-13 was completely ablated when mice were pretreated with the peripherally restricted CB_1_ antagonist AM6545, suggesting that CB-13 achieves anti-allodynia via peripheral CB_1_ receptors. This suggests that, although central mechanisms of cannabinoid-induced analgesia may differ, acute peripheral mechanisms may be independent of sex. It would be interesting to evaluate the potential sex difference of analgesic tolerance following repeated dosing with a peripherally selective CB_1_ agonist to better understand the interplay between sex and cannabinoid-mediated analgesia.

Compared to the wealth of behavioral studies linking CB_1_ receptor activation with antinociception and analgesia, relatively little is known about CB_1_ receptors and inflammation-induced changes in sensory neuron physiology. Prior studies have shown that CB_1_ receptor activation can block TRPV1 sensitization induced by NGF or bradykinin [46,57,75]. Consistent with this body of work, our calcium imaging results indicate that CB-13 reduces PGE2-induced TRPV1 sensitization in mouse DRG neurons (Figure 4). Interestingly, one recent abstract reported no effect of CB-13 on PGE2-induced TRPV1 sensitization in human sensory neurons [15]. The apparent discrepancy may be due to biological differences across species between mouse DRG and human and rat DRG, such as the lack of G protein-coupled inwardly rectifying potassium channels in mouse DRG neurons [56]. Regardless of this species difference, this work may be informative regarding the clinical utility of peripheral cannabinoid agonists in the context of different inflammatory states. Further work is needed to determine whether the effect of CB-13 generalizes to other inflammatory mediators, in both mouse and human sensory neurons. In our electrophysiology experiments, CB-13 partially attenuated PGE2-induced hyperexcitability in DRG neurons by reducing action potential firing (Figure 5). Although it has previously been reported that the endogenous cannabinoid anandamide (AEA) reduced action potential firing at baseline in rat DRG, CB-13 did not affect action potential firing in DRG neurons. These different results may be attributed to AEA metabolites interacting with voltage-gated potassium channels [23], differences in culture conditions (e.g. 1-2 days employed here vs. 7 days in culture [23]) and species differences in DRG physiology. Our findings that CB-13 reduces PGE2-induced TRPV1 sensitization and neuronal hyperexcitability support a potential analgesic role for CB_1_ activation in the periphery, and provide a potential mechanistic basis for CB-13’s anti-allodynic effects in the context of an inflammatory state.

The present study shows multiple phenotypic demonstrations of central penetrance by CB-13, including cannabinoid triad behaviors and CB_1_-mediated dependence. First, we found that acute dosing with CB-13 (10 mg/kg i.p.) produces catalepsy and tail-flick antinociception, behaviors indicative of central CB_1_ engagement. This was surprising, as the initial study of CB-13 reported that administration of CB-13 at 0.2 or 2 mg/kg p.o. did not produce catalepsy in rats [22]. However, another group later reported a slight reduction in locomotor activity at CB-13 dose of 1 mg/kg i.p. in C57BL/6N mice [8] and demonstrated that CB-13 produced hypothermia at 10 mg/kg i.v. [61] and 5 mg/kg i.p.[60] in ABH mice. The relatively smaller hypothermic effect observed here (Figure 3G-I) could represent a difference in mouse strain (ABH vs.C57BL/6J)[58,61]. Interestingly, in our study tolerance developed differently between behaviors, as 10 mg/kg of CB-13 continued to produce catalepsy but not tail-flick antinociception on day 3 of dosing (Figure 3B,E). This may reflect a difference in dose sensitivity of the behaviors induced by central CB_1_ receptor engagement, as a recent report using the peripherally restricted agonist PrNMI demonstrated alterations in the open field assay with no effect on body temperature or hot plate response[83]. Indeed, it was previously demonstrated that the ED_50_ for producing catalepsy was lower than that of tail-flick antinociception and hypothermia for the centrally penetrant cannabinoid receptor agonists CP47,497 and Δ ^9-^THC[27].

Our study is also the first to evaluate the occurrence of CB_1_ receptor dependence, a centrally mediated phenomenon [19,36,64,70,73,74], following repeated dosing with a peripherally restricted cannabinoid receptor agonist. Strikingly, rimonabant challenge following treatment with CB-13 (3 and 10 mg/kg i.p.) once daily for 9 days resulted in an increase in paw tremors and relative decrease in rimonabant-evoked scratching behaviors, indicating CB_1_ receptor dependence was induced by chronic CB-13 treatment (Figure 2B-C; Figure 3J-K). Our finding was observed in both CFA-treated and CFA-naïve mice, suggesting the presence of an inflammatory state is not required for CB-13 to penetrate the CNS. This is important, as inflammatory status can alter CB_1_ receptor density and function[49] and blood brain barrier permeability in some cases [6].

Anti-allodynic tolerance has been demonstrated following repeated dosing of centrally penetrant CB_1_ agonists such as Δ^9-^THC [19] and WIN55,212-2 [71], as well as monoacylglycerol lipase inhibitors [65,69]. Conversely, peripherally restricted cannabinoid ligands such as CB-13 and PrNMI have demonstrated no tolerance with repeated dosing in a rat models of neuropathic pain [22,52], suggesting peripherally selective cannabinoid ligands may retain anti-allodynic efficacy without the development of tolerance. Once-daily dosing with CB-13 produced anti-allodynic efficacy on days 1 and 3; surprisingly, tolerance developed by day 7 of dosing in all doses tested—including those that did not produce centrally mediated triad behaviors. This observation may reveal multiple possible interactions between CB-13 and peripheral and central nervous system components. First, these results may suggest that peripheral CB_1_ receptors may contribute to the development of anti-allodynic tolerance, in contrast to previous reports demonstrating tolerance only develops with central CB_1_ receptor agonists. Second, CB-13 also exhibits affinity for CB_2_ receptors [22], and, although CB_2_ receptor agonists demonstrate anti-allodynic efficacy that is devoid of tolerance [19,29,62,63,76,82], it may also be possible that the interplay between CB_1_ and CB_2_ receptors contributes to the development of tolerance induced by CB-13. Finally, CNS exposure to CB-13 could increase following repeated dosing. As CB-13 is postulated to be excluded from the CNS via an ATP-binding cassette transport pump, accumulation of CB-13 may result in a saturation of this pump, leading to increased penetrability into the CNS [61]. It is unclear how CNS penetrance of CB-13 would lead to anti-allodynic tolerance, as the anti-allodynia caused by CB-13 was shown to be peripheral CB_1_-mediated acutely. One interpretation may be that functional CB_1_ desensitization in the central tissues renders peripheral CB_1_ activation to become no longer effective as witnessed with other centrally penetrant cannabinoid agonists [18,19,40,50,54,71] and upregulation of 2-arachidonoylglycerol via monoacylglycerol lipase inhibition [14,39,65]. Further work is needed to better understand the mechanisms underlying anti-allodynic tolerance following repeated activation of CB_1_ receptors.

Overall, our findings show that acute treatment with a high analgesic dose of CB-13 engages central CB_1_ receptors. Although lower doses of CB-13 yielded anti-allodynia while avoiding producing central CB_1_-mediated behaviors acutely, repeated dosing lead to central CB_1_ activation in mice. Clinical reports demonstrate that CB-13 produces side effects consistent with engagement of central CB_1_ receptors (e.g. sedation, psychoactivity) at higher doses [25,38]. Another peripherally restricted CB_1_-preferring agonist, AZD1940, also produced CNS-mediated effects of sedation and “high” feeling in healthy male volunteers that were greater at a higher dose (800 µg, administered orally)[38]. Our study demonstrates how central CB_1_ activation can arise from acute high doses or repeated low doses to produce unwanted side effects and thus has important implications for the use of existing peripherally restricted CB_1_-preferring ligands. Future efforts to target peripheral CB_1_ receptors for analgesia should therefore focus on improving the pharmacokinetic properties to limit CNS penetrance even at high doses or with long-term, repeated administration.

Together, our experiments suggest that CB_1_ receptor activation in peripheral sensory neurons may lead to analgesia in the context of inflammation by reducing TRPV1 sensitization and membrane hyperexcitability induced by inflammatory mediators. Our findings also show that whereas CB-13 acutely produces peripheral CB_1_-mediated anti-allodynia, repeated dosing with CB-13 leads to anti-allodynic tolerance and central CB_1_-mediated dependence. These studies provide insight into the mechanistic basis and potential concerns of using currently available peripheral CB_1_ agonists for the treatment of chronic pain. It will be important for future studies to determine whether new molecules can be developed that maintain peripheral CB_1_ activity while reducing the potential for CNS exposure. A drug with these properties would enable clinical studies that can test the potential analgesic efficacy of full activation of peripheral CB_1_ receptors without dose-limiting side effects due to engagement of central CB_1_ receptors, and thus help improve the evidence base supporting (or not) the use of CB_1_ targeting medications for the treatment of pain[1].

## Acknowledgments

We thank the entire Gereau lab for helpful comments and critiques.

