## Supplemental tables for "The cannabinoid agonist CB-13 produces peripherally mediated analgesia in mice but elicits tolerance and signs of CNS activity with repeated dosing"

**Table 1. Statistics from Figure 1**

| **Figure 1E - Timecourse** | | | |
|---|---|---|---|
| **Type of test** | **Comparison** | **P-value summary** | **F, P-value** |
| Two-way ANOVA (BL vs post-CFA) | Time | **** | $F_{(1, 25)}$ = 238.5, P<0.0001 |
| | Treatment | ns | $F_{(3, 25)}$ = 0.2273, P=0.8765 |
| | Interaction | ns | $F_{(3, 25)}$ = 0.08162, P=0.9694 |
| **Type of test** | **Comparison** | **P-value summary** | **F, P-value** |
| Two-way ANOVA (post-CFA all timepoints) | Time | **** | $F_{(4.279, 107.0)}$ = 39.91, P<0.0001 |
| | Treatment | **** | $F_{(3, 25)}$ = 39.49, P<0.0001 |
| | Interaction | **** | $F_{(18, 150)}$ = 5.391, P<0.0001 |
| Tukey's post-hoc | | | |
| 0.5h | Vehicle vs. CB-13 (3) | **** | <0.0001 |
| | Vehicle vs. CB-13 (10) | *** | 0.0005 |
| | Vehicle vs. CB-13 (1) | *** | 0.0007 |
| | CB-13(3) vs. CB-13 (10) | ns | >0.9999 |
| | CB-13(3) vs. CB-13 (1) | ns | 0.0979 |
| | CB-13 (10) vs. CB-13 (1) | ns | 0.1977 |
| 1h | Vehicle vs. CB-13 (3) | ** | 0.0011 |
| | Vehicle vs. CB-13 (10) | *** | 0.0004 |
| | Vehicle vs. CB-13 (1) | * | 0.0251 |
| | CB-13(3) vs. CB-13 (10) | ns | 0.4816 |
| | CB-13(3) vs. CB-13 (1) | ns | 0.3266 |
| | CB-13 (10) vs. CB-13 (1) | ns | 0.061 |
| 1.5h | Vehicle vs. CB-13 (3) | * | 0.0109 |
| | Vehicle vs. CB-13 (10) | ** | 0.0033 |
| | Vehicle vs. CB-13 (1) | * | 0.011 |
| | CB-13(3) vs. CB-13 (10) | ns | 0.9759 |
| | CB-13(3) vs. CB-13 (1) | ns | 0.9091 |
| | CB-13 (10) vs. CB-13 (1) | ns | 0.6693 |
| 6h | Vehicle vs. CB-13(3) | * | 0.0128 |
| | Vehicle vs. CB-13 (10) | *** | 0.0005 |
| | Vehicle vs. CB-13 (1) | * | 0.0169 |
| | CB-13(3) vs. CB-13 (10) | ns | 0.7176 |
| | CB-13(3) vs. CB-13 (1) | ns | 0.6111 |
| | CB-13 (10) vs. CB-13 (1) | ns | 0.069 |
| 7.5h | Vehicle vs. CB-13(3) | ns | 0.0766 |
| | Vehicle vs. CB-13 (10) | **** | <0.0001 |
| | Vehicle vs. CB-13 (1) | * | 0.0398 |
| | CB-13(3) vs. CB-13 (10) | * | 0.0143 |
| | CB-13(3) vs. CB-13 (1) | ns | 0.8545 |
| | CB-13 (10) vs. CB-13 (1) | **** | <0.0001 |
| **Figure 1F** | | | |
| **Type of test** | **Comparison** | **P-value summary** | **F, P-value** |
| One-way ANOVA | Post-drug | **** | $F_{(3, 24)}$ = 4.703, P<0.0001 |
| Tukey's post-hoc | Vehicle vs. CB-13 | * | 0.0207 |
| | Vehicle vs. CB13+AM6545 | ns | 0.9989 |
| | Vehicle vs. AM6545 | ns | 0.9954 |
| | CB-13 vs. CB13+AM6545 | * | 0.0291 |
| | CB-13 vs. AM6545 | * | 0.0471 |

| | | | |
|---|---|---|---|
| | CB13+AM6545 vs. AM6545 | ns | 0.9997 |

**Table 2. Statistics from Figure 2**

| **Figure 2A - Chronic dosing** | | | |
|---|---|---|---|
| **Type of test** | **Comparison** | **P-value summary** | **F, P-value** |
| Two-way ANOVA | Time | **** | $F_{(2.388, 57.32)} = 38.29$, p<0.0001 |
| | Treatment | **** | $F_{(3, 24)} = 26.24$,p<0.0001 |
| | Interaction | **** | $F_{(9, 72)} =7.792$, p<0.0001 |
| Tukey's post-hoc | | | |
| Day 1 | Vehicle vs. CB-13(3) | *** | 0.0002 |
| | Vehicle vs. CB-13 (10) | *** | 0.0005 |
| | Vehicle vs. CB-13 (1) | ** | 0.0011 |
| | CB-13(3) vs. CB-13 (10) | ns | 0.9921 |
| | CB-13(3) vs. CB-13 (1) | ns | 0.3308 |
| | CB-13 (10) vs. CB-13 (1) | ns | 0.2859 |
| Day 3 | Vehicle vs. CB-13(3) | **** | <0.0001 |
| | Vehicle vs. CB-13 (10) | ** | 0.0016 |
| | Vehicle vs. CB-13 (1) | * | 0.0486 |
| | CB-13(3) vs. CB-13 (10) | ns | 0.9981 |
| | CB-13(3) vs. CB-13 (1) | * | 0.0206 |
| | CB-13 (10) vs. CB-13 (1) | ns | 0.0931 |
| **Withdrawal** | | | |
| **Figure 2B - Paw Tremor** | | | |
| **Type of test** | **Comparison** | **P-value summary** | **F, P-value** |
| One-Way ANOVA | All | ** | $F_{(3, 24)} = 5.971$,p=0.0034 |
| Tukey's post-hoc | Vehicle vs. CB-13 (1) | ns | 0.9499 |
| | Vehicle vs. CB-13 (3) | ns | 0.2965 |
| | Vehicle vs. CB-13 (10) | ** | 0.0037 |
| | CB-13 (1) vs. CB-13 (3) | ns | 0.5906 |
| | CB-13 (1) vs. CB-13 (10) | * | 0.0132 |
| | CB-13 (3) vs. CB-13 (10) | ns | 0.1872 |
| **Figure 2C- Scratching** | | | |
| **Type of test** | **Comparison** | **P-value summary** | **F, P-value** |
| One-way ANOVA | All | *** | $F_{(3, 24)} = 10.78$, P=0.0001 |
| Tukey's post-hoc | Vehicle vs. CB-13 (1) | ns | 0.302 |
| | Vehicle vs. CB-13 (3) | ** | 0.0048 |
| | Vehicle vs. CB-13 (10) | *** | 0.0001 |
| | CB-13 (1) vs. CB-13 (3) | ns | 0.2207 |
| | CB-13 (1) vs. CB-13 (10) | ** | 0.0086 |
| | CB-13 (3) vs. CB-13 (10) | ns | 0.4261 |

**Table 3. Statistics from Figure 3**

| **Catalepsy** | | | |
|---|---|---|---|
| **Figure 3A - Day 1** | | | |
| **Type of test** | **Comparison** | **P-value summary** | **F, P-value** |
| Two-way ANOVA | Treatment | ** | $F_{(2, 16)}$ = 11.63, P=0.008 |
| | Time | **** | $F_{(2.579, 41.26)}$ =10.79, P<0.0001 |
| | Interaction | **** | $F_{(14, 112)}$ = 8.983, P<0.0001 |
| Tukey's post-hoc | | | |
| 6h | CB-13(3) vs. CB-13 (10) | ns | 0.052 |
| | CB-13(3) vs. Vehicle | ns | 0.3487 |
| | CB-13 (10) vs. Vehicle | * | 0.0378 |
| 7.5h | CB-13(3) vs. CB-13 (10) | * | 0.0268 |
| | CB-13(3) vs. Vehicle | ns | 0.5084 |
| | CB-13 (10) vs. Vehicle | * | 0.0214 |
| **Figure 3B - Day 3** | | | |
| **Type of test** | **Comparison** | **P-value summary** | **F, P-value** |
| Two-Way ANOVA | Treatment | * | $F_{(2, 16)}$ =4.267, P=0.0327 |
| | Time | *** | $F_{(1.668, 26.68)}$ = 10.86, P=0.0007 |
| | Interaction | *** | $F_{(14, 112)}$ = 3.347, P=0.0002 |
| Tukey's post-hoc | | | |
| 3h | CB-13 (3) vs. CB-13 (10) | ns | 0.14 |
| | CB-13 (3) vs. Vehicle | ns | 0.2698 |
| | CB-13 (10) vs. Vehicle | * | 0.0454 |
| **Figure 3C - Day 7** | | | |
| **Type of test** | **Comparison** | **P-value summary** | **F, P-value** |
| Two-way ANOVA | Treatment | Ns | $F_{(2, 16)}$ = 3.200, P=0.0678 |
| | Time | *** | $F_{(1.863, 29.80)}$ =11.67, P=0.0002 |
| | Interaction | *** | $F_{(14, 112)}$ = 3.210, P=0.0003 |
| **Tail-Flick** | | | |
| **Figure 3D - Day 1** | | | |
| **Type of test** | **Comparison** | **P-value summary** | **F, P-value** |
| Two-way ANOVA | Treatment | ** | $F_{(2, 16)}$ = 6.422, P=0.0090 |
| | Time | **** | $F_{(3.050, 48.80)}$ = 14.39, P<0.0001 |
| | Interaction | **** | $F_{(14, 112)}$ = 5.921, P<0.0001 |
| Tukey's post-hoc | | | |
| 3h | CB-13 (3) vs. CB-13 (10) | * | 0.0231 |
| | CB-13 (3) vs. Vehicle | ns | 0.926 |
| | CB-13 (10) vs. Vehicle | * | 0.0277 |
| **Figure 3E – Day 3** | | | |
| **Type of test** | **Comparison** | **P-value summary** | **F, P-value** |
| Two-way ANOVA | Treatment | ns | $F_{(2, 16)}$ = 1.603, P=0.2319 |
| | Time | * | $F_{(3.226, 51.62)}$ = 2.746, P=0.0485 |
| | Interaction | ns | $F_{(14, 112)}$ = 1.357, P=0.1862 |
| **Figure 3F – Day 7** | | | |
| **Type of test** | **Comparison** | **P-value summary** | **F, P-value** |
| Two-way ANOVA | Treatment | ns | $F_{(2, 16)}$ = 1.791, P=0.1986 |
| | Time | * | $F_{(2.707, 43.31)}$ = 3.756, P=0.0206 |
| | Interaction | ns | $F_{(14, 112)}$ = 1.416, P=0.1571 |
| **Body Temperature** | | | |
| **Figure 3G – Day 1** | | | |
| **Type of test** | **Comparison** | **P-value summary** | **F, P-value** |
| Two-way ANOVA | Treatment | ns | $F_{(2, 16)}$ = 1.948, P=0.1749 |
| | Time | ** | $F_{(2.017, 32.28)}$ = 6.075, P=0.0027 |
| | Interaction | ** | $F_{(14, 112)}$ = 2.255, P=0.0096 |

| **Figure 3H – Day 3** | | | |
|---|---|---|---|
| **Type of test** | **Comparison** | **P-value summary** | **F, P-value** |
| Two-way ANOVA | Treatment | ns | $F_{(2, 16)}$ = 3.546, P=0.0531 |
| | Time | ** | $F_{(4.864, 77.82)}$ = 4.078, P=0.0027 |
| | Interaction | ns | $F_{(14, 112)}$ = 0.5532, P=0.8953 |
| **Figure 3I – Day 7** | | | |
| **Type of test** | **Comparison** | **P-value summary** | **F, P-value** |
| Two-way ANOVA | Treatment | ns | $F_{(2, 16)}$ = 0.7193, P=0.5022 |
| | Time | ** | $F_{(4.007, 64.12)}$ = 4.878, P=0.0017 |
| | Interaction | * | $F_{(14, 112)}$ = 2.122, P=0.0155 |
| **Withdrawal** | | | |
| **Figure 3J-Paw Tremor** | | | |
| **Type of Test** | **Comparison** | **P-value summary** | **F, P-value** |
| One-way ANOVA | All | * | $F_{(12,15)}$ = 4.637, P=0.0271 |
| Tukey's post-hoc | Vehicle vs. CB-13(3) | ns | 0.6369 |
| | Vehicle vs. CB-13(10) | * | 0.0242 |
| | CB-13(3) vs. CB-13(10) | ns | 0.1335 |
| **Figure 3K- Scratching** | | | |
| **Type of Test** | **Comparison** | **P-value summary** | **F, P-value** |
| One-way ANOVA | All | **** | $F_{(2,15)}$ = 40.94, P<0.0001 |
| Tukey's post-hoc | Vehicle vs. CB-13(3) | **** | <0.0001 |
| | Vehicle vs. CB-13(10) | **** | <0.0001 |
| | CB-13(3) vs. CB-13(10) | ns | 0.2025 |

**Table 4. Statistics from Figure 4**

| **Figure 4A - Hargreaves** | | | |
|---|---|---|---|
| **Type of test** | **Comparison** | **P-value summary** | **F, P-value** |
| Two-way ANOVA (BL vs. Post-CFA) | Treatment | ns | $F_{(1, 28)}$ = 0.1716, P=0.6818 |
| | Time | ******** | $F_{(1, 28)}$ = 109.1, P<0.0001 |
| | Interaction | ns | $F_{(1, 28)}$ = 0.1945,P=0.6626 |
| **Type of test** | **Comparison** | **P-value summary** | **F, P-value** |
| Two-way ANOVA (Post-CFA all timepoints) | Treatment | ***** | $F_{(1, 28)}$ = 6.331,P=0.0179 |
| | Time | ******** | $F_{(4.674, 130.9)}$ = 9.157, P<0.0001 |
| | Interaction | ***** | $F_{(7, 196)}$ = 2.563, P=0.0151 |
| Tukey's post-hoc | | | |
| 5h | Vehicle vs. CB-13 (3) | * | 0.042 |
| **Figure 4E - Calcium imaging** | | | |
| **Type of test** | **Comparison** | **P-value summary** | **F, P-value** |
| One-way ANOVA | All | ****** | $F_{(2, 139)}$ = 7.074, P=0.0012 |
| Tukey's post-hoc | Cap vs. PGE2 | * | 0.0176 |
| | Cap vs. PGE2+CB13 | ns | 0.5393 |
| | PGE2 vs. PGE2+CB13 | ** | 0.0026 |

**Table 5. Statistics from Figure 5**

| **Figure 5A** | | | |
|---|---|---|---|
| **Type of test** | **Fixed effects** | **P-value summary** | **F, P-value** |
| Mixed effects model | Current step | **** | $F_{(1.988, 19.88)}$ = 21.50, P<0.0001 |
| | PGE2 | ** | $F_{(1.000, 10.00)}$ = 11.82, P=0.0063 |
| | Interaction | ns | $F_{(1.972, 14.46)}$ = 1.049, P=0.3746 |
| Post-hoc tests (Sidak's) | **Current step** | **P-value summary** | **P-value** |
| Pre- vs. Post-PGE2 | 1x | ns | 0.1069 |
| | 2x | ns | 0.0598 |
| | 3x | ns | 0.3076 |
| | 4x | ns | 0.0806 |
| **Figure 5B** | | | |
| **Type of test** | **Fixed effects** | **P-value summary** | **F, P-value** |
| Mixed effects model | Current step | **** | $F_{(1.019, 10.19)}$ = 46.48, P<0.0001 |
| | CB-13+PGE2 | ns | $F_{(1.000, 10.00)}$ = 0.9501, P=0.3527 |
| | Interaction | ns | $F_{(1.459, 11.19)}$ = 0.4223, P=0.604 |
| **Figure 5C** | | | |
| **Type of test** | **Fixed effects** | **P-value summary** | **F, P-value** |
| Mixed effects model | Current step | **** | $F_{(1.542, 15.42)}$ = 21.93, P<0.0001 |
| | CB-13 | ns | $F_{(1.000, 10.00)}$ = 1.184, P=0.3021 |
| | Interaction | ns | $F_{(1.691, 16.35)}$ = 0.9325, P=0.3986 |
| **Figure 5D** | | | |
| **Type of test** | **Fixed effects** | **P-value summary** | **F, P-value** |
| Mixed effects model | Current step | **** | $F_{(2.004, 20.04)}$ = 25.52, P<0.0001 |
| | Vehicle | ns | $F_{(1.000, 10.00)}$ = 0.7505. P=0.4066 |
| | Interaction | ns | $F_{(1.938, 19.38)}$ = 0.7639, P=0.4755 |
| **Figure 5E** | | | |
| **Type of test** | **Fixed effects** | **P-value summary** | **F, P-value** |
| Mixed effects model (REML) | Current step | **** | $F_{(2.243, 77.77)}$ = 87.85, P<0.0001 |
| | Treatment | ** | $F_{(3, 40)}$ = 4.535, P=0.0079 |
| | Interaction | ns | $F_{(9, 104)}$ = 1.941, P=0.054 |
| *Tukey's post-hoc* | *Comparison* | *P-value summary* | *P-value* |
| 4x current step | Vehicle vs. CB-13 | ns | 0.8848 |
| | Vehicle vs. PGE2 | * | 0.0309 |
| | Vehicle vs. CB13+PGE2 | ns | 0.9399 |
| | CB-13 vs. PGE2 | * | 0.0141 |
| | CB-13 vs. CB13+PGE2 | ns | 0.5586 |
| | PGE2 vs. CB13+PGE2 | ns | 0.0766 |
| **Figure 5F** | | | |
| **Type of test** | **Comparison** | **P-value summary** | **F, P-value** |
| One way ANOVA | All | *** | $F_{(3, 40)}$ = 7.534, P=0.0004 |
| Tukey's post-hoc | $PGE_2$ vs. CB13 + $PGE_2$ | ns | 0.8541 |
| | $PGE_2$ vs. CB-13 | ns | 0.1883 |
| | $PGE_2$ vs. Vehicle | ** | 0.0067 |

| | | | |
|---|---|---|---|
| | CB13 + $PGE_2$ vs. CB-13 | * | 0.0337 |
| | CB13 + $PGE_2$ vs. Vehicle | *** | 0.0007 |
| | CB-13 vs. Vehicle | ns | 0.4937 |
| **Figure 5G** | | | |
| **Type of test** | **Comparison** | **P-value summary** | **F, P-value** |
| Two-way ANOVA | Time | ns | $F_{(1.478, 57.66)} = 2.341$, P=0.1192 |
| | Treatment | ns | $F_{(6, 78)} = 0.6003$, P=0.7292 |
| | Interaction | ns | $F_{(3, 39)} = 1.147$, P=0.3423 |
| **Figure 5H** | | | |
| **Type of test** | **Comparison** | **P-value summary** | **F, P-value** |
| One-way ANOVA | All | ns | $F_{(3, 40)} = 1.586$, P=0.2077 |
| **Figure 5I** | | | |
| **Type of test** | **P-value summary** | **H Statistic** | **P-value** |
| Kruskal-Wallis | ns | 4.336 | 0.2274 |
